# Testing for gene-environment (GxE) interaction using p-value aggregation identifies many GxE loci

**DOI:** 10.64898/2026.02.24.707798

**Authors:** Saurabh Mishra, Rima Rati Patra, Ananya S Reddy, Abhijit Mandal, Arunabha Majumdar

## Abstract

Genome-wide gene-environment (GxE) interaction studies have seen limited success in detecting reliable GxE signals. A standard genome-wide GxE scan assumes a single genetic mode of inheritance, such as an additive model. It can lead to reduced statistical power when the true genetic model is non-additive, such as a recessive model. We propose a robust GxE testing approach that uses Cauchy p-value aggregation. It combines the p-values from GxE tests based on the additive, dominant, and recessive genetic models. Using extensive simulation studies, we demonstrate that the p-value combination strategy offers a robust and powerful approach to identifying GxE interactions regardless of the underlying genetic model. The method is substantially more powerful than the additive model when the true genetic model is recessive. It is also more powerful than the general two-degree-of-freedom genotypic test for GxE interaction. We apply our approach to analyze GxE interactions in the UK Biobank data across several combinations of phenotypes and environmental factors. For glycated hemoglobin (HbA1c) level, treating cumulative smoking exposure as the lifestyle factor, our approach identified 82 independent GxE loci while controlling FDR at 5%. The GxE test based on the additive genetic model detected 24 loci. For type 2 diabetes with sleep duration as a lifestyle factor, the proposed approach detected 563 independent GxE loci at 5% FDR, substantially exceeding the number of discoveries by the other approaches.

## 1 Introduction

Interactions between genetic and environmental factors (GxE) impact the risk of complex human diseases [1]. Analyzing GxE interactions is crucial for understanding how environmental factors modulate genetic predisposition to a complex phenotype [2]. Genome-wide association studies (GWAS) have become a standard approach for identifying genetic loci associated with complex phenotypes. However, GxE interaction studies have seen limited success, mainly due to inadequate statistical power [3]. A contributor to this power deficit, which is often overlooked, is the problem of incorrect genotype coding. In a standard GWAS, we assume a specific genetic mode of inheritance while testing for an association and keep it fixed for all SNPs. The genetic model describes how alleles at a marker locus combine to influence a phenotype. The most commonly used genetic models are the additive, dominant, and recessive modes of inheritance. However, the true mode of inheritance is never known *a priori*. The common practice is to assume an additive model for genotype coding for all SNPs. This can lead to power loss if the true genetic model is non-additive [4], such as a dominant or recessive model. Recent works [5] have demonstrated that a majority of complex phenotype heritability is additive heritability. However, GxE heritability is limited for complex phenotypes [6], mainly because GxE interaction effects are minor. Thus, a misspecification of the genetic model can lead to power loss while searching for genome-wide SNP-level GxE effects.

Several strategies have been developed to mitigate misspecification in the genetic model. In a genomewide GxE study, the two degrees of freedom (2df) GxE test offers a robust, genetic-model-free approach [7]. Instead of choosing a genotype coding, this method treats genotypes as categories and thus implements a 2df test. We refer to this genotype-level GxE testing approach as the 2df test. Moore et al. [7] studied power and sample size calculations for the 2df GxE test. While this framework clearly guides a study design, to our knowledge, it has not yet been implemented and systematically evaluated for a primary genome-wide GxE scan in large-scale biobank data. This creates an opportunity to both operationalize the 2df GxE test at scale and to compare it directly with model-specific and p-value aggregation strategies for robust GxE analysis. Alternatively, we can perform a GxE interaction test separately for the three genotype coding schemes. Then, we can implement a multiple testing correction, such as the Bonferroni or Holm’s method, to adjust for multiple comparisons. However, such corrections make the testing procedure conservative. A more sophisticated approach, the MAX3 test, which takes the maximum of three test statistics, relies on computationally intensive resampling procedures to derive a valid p-value. Such a resampling procedure is challenging to implement for a genome-wide scan [8, 9], mainly due to the stringent genome-wide threshold of p-values, in the order of 10*^-^*^6^, 10*^-^*^7^, 10*^-^*^8^.

In recent literature, p-value aggregation across multiple models has become a useful and popular approach. Such a strategy combines signals from one or more models to build a robust and powerful approach. Two well-known p-value combination approaches are the Harmonic mean aggregation [10], and the Cauchy aggregation [11, 12]. The main advantage of these methods is that the combined p-value obtained from model averaging is well-calibrated for error control in a multiple-comparison set-up. At the same time, these methods are computationally very fast, making them easily implementable at a genome-wide scale.

In this article, we propose using p-value aggregation to address the uncertainty in selecting the correct genotype coding while performing a GxE test. We obtain three different p-values for testing a GxE interaction, based on additive, dominant, and recessive genetic modes of inheritance. Next, we implement p-value aggregation to obtain a single p-value for testing GxE interaction while accounting for the uncertainty in the true genetic mode of inheritance. We refer to this procedure as the GETAP (**G**x**E T**esting using **A**ggregated **P**-value) approach. We hypothesize that combining statistical evidence from GxE analyses based on multiple, distinct genetic models via p-value aggregation can construct a single, powerful, and robust testing procedure that is sensitive to a GxE signal regardless of the true underlying genetic model. Suppose for a GxE interaction between a SNP and an environmental factor, the underlying genetic mode of inheritance is recessive. According to common practice, an additive genetic model is incorrect and can lead to power loss. However, the recessive model-based GxE interaction p-value is expected to be strong. P-value aggregation of the multiple models is likely to retain and reflect the signal. The Cauchy combination of multiple p-values offers a valid combined p-value under general dependence of the tests, which is crucial for such an aggregation approach [11, 12].

Using extensive simulations, we compare the GETAP approach with the individual additive, dominant, and recessive genetic models, as well as the 2df GxE testing approach, with respect to type I error rate and power. We then apply these approaches to several phenotype-environment combinations in the UK Biobank (UKB). The quantitative phenotypes include glycated hemoglobin (HbA1c), pulmonary function (FEV_1_/FVC), body mass index (BMI), C-reactive protein (CRP), and triglycerides. The environmental exposures include cumulative smoking exposure (pack-years), smoking status, sleep duration, alcohol intake frequency, and healthy diet score. In addition, we analyze two binary disease outcomes, type 2 diabetes and chronic obstructive pulmonary disease (COPD), considering sleep duration and cumulative smoking exposure as the lifestyle factors, respectively.

## 2 Methods

We consider the generalized linear model (GLM) as the main framework for testing GxE interactions. For an individual *i*, let *Y_i_* be the phenotype, *G_T,i_* be the correctly coded numeric genotype value, *E_i_* be the environmental exposure, and ***C****_i_* be a vector of adjusting covariates, *i* = 1*, …, n*. ***C****_i_* includes age, sex, principal components (PCs) of genetic ancestry, etc. We have the following GLM:

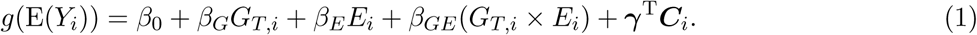

Here, *g*(*·*) is the link function, *β_G_* is the marginal main genetic effect of a SNP on the phenotype, *β_E_* is the marginal exposure effect, and *β_GE_* is the GxE interaction effect on the phenotype. γ denotes the vector of effect sizes for the adjusting covariates. We can choose the link function depending on the phenotype type; for example, an identity link function for a continuous quantitative phenotype or a logit link function for a binary disease phenotype. We assume that *G_T,i_* is the ideal genotype coding, i.e., one of the additive, dominant, or recessive coding. Here, the effect sizes are defined for the true genetic model. To evaluate the GxE interaction, we test for *H*_0_: *β_GE_* = 0 vs. *β_GE_ ≠* 0. For a normally distributed quantitative phenotype, we consider:

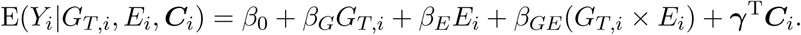

For a binary disease phenotype:

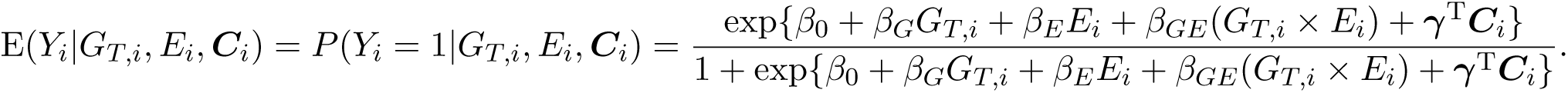

In reality, *G_T,i_*is unknown. We denote the chosen model for genotype coding as *G_M_*, where *M* = *A* (additive), *D* (dominant), and *R* (recessive) (Table 1). Given a choice of the genetic mode of inheritance *M* = *A, D, R*, we model the relationship between the phenotype and the genotype and the environmental factor as:

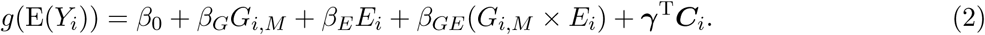

We test for the GxE interaction by implementing the one-degree-of-freedom test based on this model. For *M* = *A, D, R*, we repeat this test and obtain three different p-values of testing the GxE interaction.

**Table 1:** Genotype coding schemes for additive (A), dominant (D), recessive (R), and 2df models. *A* denotes the minor/risk allele, and the three possible genotypes are *aa*, *aA*, and *AA*.

| Model | $aa$ | $aA$ | $AA$ |
| --- | --- | --- | --- |
| Additive ($G_A$) | 0 | 1 | 2 |
| Dominant ($G_D$) | 0 | 1 | 1 |
| Recessive ($G_R$) | 0 | 0 | 1 |
| 2df ($G_{Het}, G_{Hom}$) | (0,0) | (1,0) | (0,1) |

### 2.1 Cauchy combination

Suppose we implement *k* different models for testing the GxE interaction and obtain *k* p-values: *p*_1_*,…, p_k_*. Given these dependent p-values obtained from the *k* different statistical tests, the aggregated Cauchy association test (ACAT) combines them through Cauchy transformation and weighted aggregation [11, 12]. The ACAT statistic is given by:

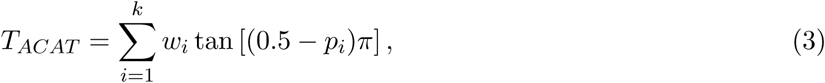

where *w_i_* ≥ 0*, i* = 1*,…, k*, are user-specified weights with 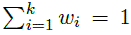. In our context, we employ uniform weights, *w_i_*= 1*/k*. We assume that no prior information prioritizing a specific genotype coding is available. The aggregated p-value is obtained from the cumulative distribution function of the standard Cauchy distribution:

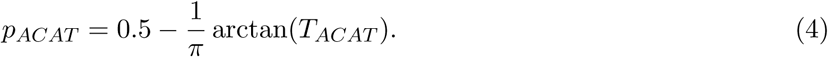

This formula provides an approximate aggregated p-value under the null hypothesis. For a given number of tests combined, the approximation is highly accurate near the tail of the null distribution [11], which is important for a genome-wide study. Here, the significant p-values are small. The approximation is reliable for arbitrary dependence between the input p-values, which is crucial in the current context. Due to these robustness properties and computational simplicity, the Cauchy combination has become a reliable alternative for p-value aggregation. For our proposed GETAP approach, we aggregate GxE testing p-values obtained from the three most-used genetic models:

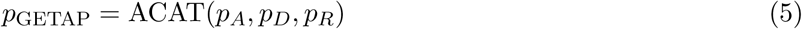

where *p_A_*, *p_D_*, and *p_R_* are the GxE interaction p-values from the additive, dominant, and recessive models, respectively, applied to the same pair of SNP-environmental factor (Figure 1). *p*_ACAT_ evaluates whether there is a signal of GxE interaction based on at least one of the three genetic models. Therefore, the global null hypothesis states that *H*_0_: *β_GE_* = 0 for all three different genetic modes of inheritance, versus *H*_1_: *β_GE_ ≠* 0 for at least one of the three genetic models.

**Figure 1:**
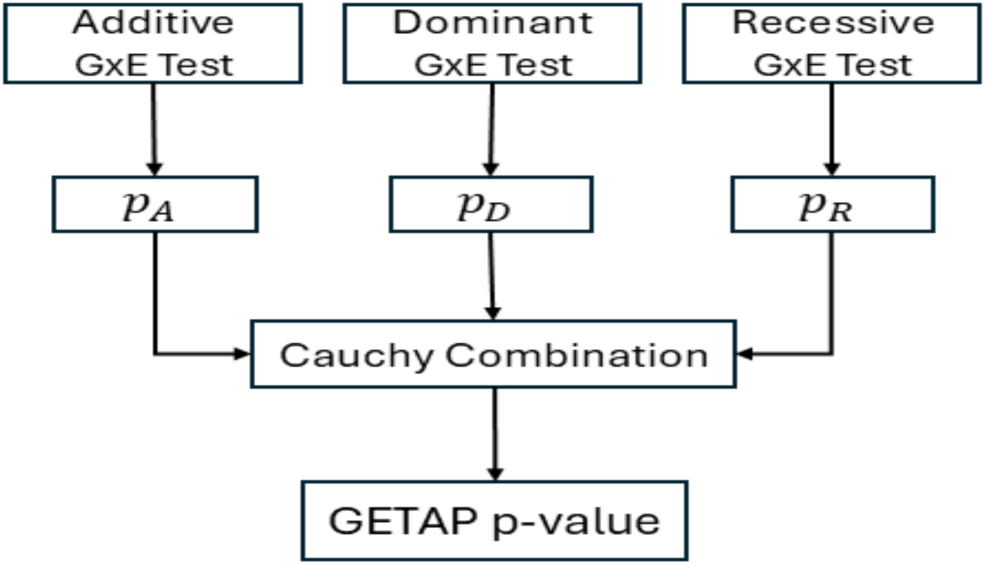
Overview of the GETAP GxE testing procedure. We combine the p-values from additive (*pA*), dominant (*pD*), and recessive (*pR*) G × E tests via Cauchy aggregation to produce the GETAP p-value.

### 2.2 P-value aggregation using the Harmonic mean p-value (HMP)

The Harmonic Mean p-value (HMP) [10] is obtained by combining multiple possibly dependent p-values into a single global significance measure. Consider *k* p-values, *p*_1_*,…, p_k_,* for testing GxE interaction based on multiple genetic models. Let *w*_1_*,…, w_k_* be nonzero positive weights satisfying 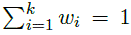. The HMP statistic is defined as:

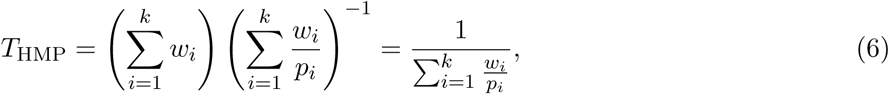

where the second equality uses 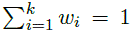. For equal weights *w_i_* = 1*/k*, this reduces to the classical harmonic mean:

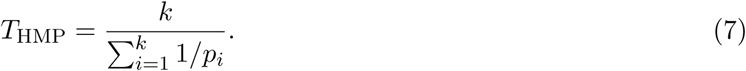

Under the global null hypothesis that there is no GxE interaction effect based on the individual genetic models, the Landau stable law is used to approximate the null distribution (more details provided in the reference paper [10]).

A key feature of the HMP procedure is that the aggregated p-value remains valid even when the original p-values are dependent. Thus, *p*_HMP_ defines an overall p-value for evaluating the global null hypothesis.

The weights *w_i_*s can be unequal, though we adopt equal weights: *w_i_* = 1*/k*. The true underlying genetic model is not known. The aggregated p-value is given by:

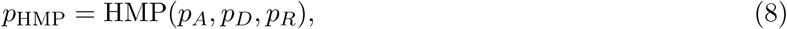

where *p_A_*, *p_D_*, and *p_R_* are the GxE test interaction p-values from the additive, dominant, and recessive models, respectively, applied to the same pair of SNP-environmental factor.

### 2.3 2 degree of freedom (2df) GxE test

The 2df GxE test uses two indicator variables, *G_Het,i_*for the heterozygous genotype and *G_Hom,i_* for the homozygous risk genotype (Table 1). It considers:

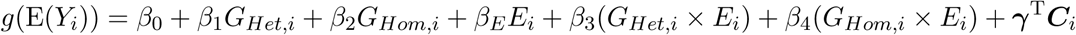

We test for the null hypothesis of no GxE interaction based on a joint test of the two interaction coefficients: *H*_0_: *β*_3_ = *β*_4_ = 0 vs. *H*_1_: *β*_3_ ≠ 0 or *β*_4_ ≠ 0 or both. Hence, this is a two-degree-of-freedom (2df) test.

## 3 Simulation study

We performed extensive simulations to evaluate the validity, robustness, and power of the proposed GETAP approach for identifying gene-environment interactions. We compared the approach with the additive, dominant, and recessive genetic models, and the 2df testing procedure in terms of the type I error rate and the power of detecting an interaction. We consider various simulation scenarios that comprise both continuous and binary phenotypes, as well as continuous and binary environmental variables.

### 3.1 Simulation design

While designing the simulation studies, we vary the minor allele frequency of a SNP and the magnitude of interaction effects. We consider three genetic models: additive, dominant, and recessive. We evaluate the performance of the various GxE testing procedures for a given true mode of genetic inheritance. Naturally, we expect the GxE test based on the true genetic model to perform best, but this is unknown in practice; it reflects the uncertainty regarding the genetic model in real-life studies. Therefore, it enables a direct assessment of the power loss incurred by model misspecification and the extent to which our proposed GETAP approach can recover this loss through p-value aggregation.

In each simulation replicate, we simulated genotypes for a biallelic SNP satisfying the Hardy-Weinberg equilibrium (HWE) with the minor allele frequency (MAF) ranging from 0.05 to 0.3. From the additive genotype count *G*_add_ ∊ *{*0, 1, 2*}*, we derived the other two genotype encodings - dominant: *G*_dom_ = 𝕀(*G*_add_ ≥ 1), and recessive: *G*_rec_ = 𝕀(*G*_add_ = 2), where I denotes the indicator function. For the 2df GxE test, we additionally constructed indicators of heterozygous and homozygous-alternative genotypes,

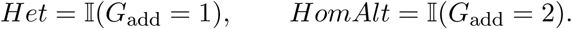

We simulate environmental exposures according to the different types of them observed in epidemiological cohorts. We generate continuous exposures from a standard normal distribution: *E_i_* ∼ *N* (0, 1). We sample binary exposures from a Bernoulli distribution with prevalence 0.2: *E_i_* ∼ Bernoulli(0.2). We simulate the phenotypes based on models that include the main genetic, main environmental, and GxE interaction effects. For continuous outcomes, we use:

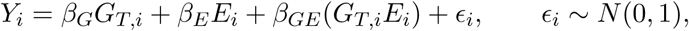

and for binary outcomes,

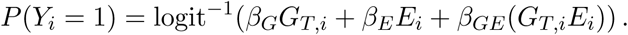

Here, *G_T,i_* denotes the encoded genotype based on the true genetic model. We fix the main genetic and environmental effects as: *β_G_*= *β_E_* = 0.2, representing a modest but realistic effect size. We vary the interaction effect *β_GE_* in a range comprising weak to strong effects: from 0.025 to 1. This allows us to assess the power of the competing approaches across different strengths of the interaction effects. We evaluate the type I error control with *β_GE_*= 0 and the above setting of the remaining parameters. We consider 10,000 individuals in the sample. For each simulation scenario, we performed 2,000 iterations. We choose ∊*_i_* ∼ *N* (0, 1) assuming that a single SNP explains a negligible proportion of complex phenotype heritability.

For each replicate in a given simulation scenario, we compute the GxE testing p-values by six different analytical strategies: three GxE tests based on three different genotype codings, p-value aggregation by Cauchy and Harmonic combination, and the general 2df GxE test. The three single-model 1df GxE tests, additive, dominant, and recessive, were implemented by fitting generalized linear models for each genotype coding and testing *H*_0_: *β_GE_*= 0 vs. *H*_1_: *β_GE_* ≠ 0 using likelihood ratio tests. We applied GETAP using the Cauchy and Harmonic p-value aggregation strategies. GETAP-ACAT combined the three p-values using the Aggregated Cauchy combination with equal weights: *w_i_* = 1*/*3. GETAP-HMP used the harmonic mean p-value with equal weights: *w_i_*= 1*/*3. For the 2df GxE test, we fit a full model including *Het*, *HomAlt*, the environmental exposure *E*, and two interaction terms *Het* × *E* and *HomAlt* × *E*, and then implemented the two-degree-of-freedom likelihood ratio test to evaluate the null hypothesis *H*_0_: *β*_3_ = *β*_4_ = 0 vs. *H*_1_: *β*_3_ ≠ 0*, β*_4_ ≠ 0, or both.

We estimated the type I error rate (TIER) for *β_GE_* = 0 at the level of significance *α* = 0.05. We computed the power at the same threshold for all non-null scenarios. To evaluate the consequences of model misspecification, we contrast the power estimates considering the analysis based on the true genetic model as the gold standard. For example, in a simulation scenario where we generated genotype data according to a dominant inheritance model, we compared the performance of the GxE test based on dominant genotype coding (correctly specified), the additive and recessive genotype codings (misspecified), the two GETAP procedures (model-averaged), and the 2df genotypic test (model-free).

### 3.2 Simulation results

We compared the GETAP strategies against the standard 1-degree-of-freedom (1df) tests assuming additive (Add), dominant (Dom), or recessive (Rec) genetic models, as well as the model-free 2df genotypic test. We evaluated the empirical Type I Error Rate (TIER) and power across a range of minor allele frequencies (MAF: 0.05, 0.1, 0.2, 0.3). We encompass various types of environmental exposures and phenotypes, such as continuous and binary.

### 3.3 Type I error rate (TIER) control

For the continuous phenotype, the various methods (Add, Dom, Rec, ACAT, HMP, 2df) demonstrate an overall adequate control of TIER (Table 2) across different simulation scenarios. Empirical TIER rates fluctuate marginally around the nominal 0.05 level, ranging between 0.04 and 0.061. This confirms the validity of all methods regarding adequate control of TIER for continuous outcomes, regardless of the underlying true genetic model, MAF, or type of environmental exposures. For the binary phenotype (Table S1), at lower and common MAFs (0.1 to 0.3), all methods exhibit controlled TIERs, with most estimates between 0.04 to 0.06. However, for MAF = 0.05, we observe some inflation in specific scenarios for the misspecified genetic-model-based 1df tests. Subsequently, the GETAP approaches had moderate inflation due to p-value aggregation. We observed the highest inflation for the recessive model for both binary and continuous environmental factors, with a TIER estimate up to 0.076 (Table S1). Such inflation is a known issue for variants with lower MAF and binary outcomes due to the smaller size of the rare homozygous genotype group. It also affected the 2df test, with a TIER estimate up to 0.065 (Table S1). On the other hand, the additive and dominant models remained better-calibrated. As a consequence, the GETAP approaches had comparatively moderate inflation: GETAP-ACAT yielded a TIER estimate up to 0.064, and HMP yielded a TIER estimate up to 0.058 (Table S1). Therefore, the TIER control pattern of the constituent tests affects the TIER control of a p-value aggregation method. However, the TIER inflation for GETAP was consistently lower than that of the 1df recessive and 2df tests, demonstrating a partial impact. In additional simulations with lower MAF (results not included to save space), we observed that TIER inflation decreased as sample sizes increased, mainly because the rare homozygous genotype group grew larger in size. Thus, the GETAP approach overall controls the type I error rate. Occasional limited fluctuations should not occur, especially in contemporary GWAS data with large sample sizes.

**Table 2:** Empirical type I error rates (TIER) for different methods of GxE interaction analyses for a continuous phenotype under possible genetic model misspecification. Results are stratified by environmental variable type (Env type: binary or continuous), minor allele frequency (0.05 - 0.3), and the true genetic model (additive, dominant, recessive). We use 2, 000 iterations in a simulation scenario to estimate the TIER. The sample comprises *n* = 10, 000 individuals.

| **True model** | **Env type** | **MAF** | **Add** | **Dom** | **Rec** | **ACAT** | **HMP** | **2df** |
| --- | --- | --- | --- | --- | --- | --- | --- | --- |
| additive | binary | 0.05 | 0.051 | 0.049 | 0.047 | 0.045 | 0.042 | 0.042 |
| additive | binary | 0.10 | 0.047 | 0.048 | 0.049 | 0.053 | 0.046 | 0.046 |
| additive | binary | 0.20 | 0.053 | 0.054 | 0.046 | 0.052 | 0.047 | 0.050 |
| additive | binary | 0.30 | 0.053 | 0.054 | 0.053 | 0.053 | 0.048 | 0.057 |
| additive | continuous | 0.05 | 0.052 | 0.049 | 0.056 | 0.057 | 0.050 | 0.061 |
| additive | continuous | 0.10 | 0.051 | 0.047 | 0.054 | 0.053 | 0.047 | 0.051 |
| additive | continuous | 0.20 | 0.045 | 0.050 | 0.047 | 0.044 | 0.040 | 0.045 |
| additive | continuous | 0.30 | 0.059 | 0.055 | 0.052 | 0.059 | 0.052 | 0.046 |
| dominant | binary | 0.05 | 0.053 | 0.046 | 0.053 | 0.059 | 0.053 | 0.059 |
| dominant | binary | 0.10 | 0.053 | 0.049 | 0.052 | 0.057 | 0.049 | 0.051 |
| dominant | binary | 0.20 | 0.052 | 0.048 | 0.053 | 0.054 | 0.048 | 0.055 |
| dominant | binary | 0.30 | 0.047 | 0.050 | 0.047 | 0.051 | 0.045 | 0.051 |
| dominant | continuous | 0.05 | 0.060 | 0.055 | 0.045 | 0.056 | 0.048 | 0.052 |
| dominant | continuous | 0.10 | 0.050 | 0.049 | 0.042 | 0.049 | 0.044 | 0.045 |
| dominant | continuous | 0.20 | 0.046 | 0.048 | 0.045 | 0.043 | 0.041 | 0.047 |
| dominant | continuous | 0.30 | 0.048 | 0.049 | 0.058 | 0.055 | 0.048 | 0.057 |
| recessive | binary | 0.05 | 0.048 | 0.045 | 0.049 | 0.048 | 0.042 | 0.044 |
| recessive | binary | 0.10 | 0.053 | 0.056 | 0.048 | 0.052 | 0.047 | 0.049 |
| recessive | binary | 0.20 | 0.040 | 0.044 | 0.047 | 0.039 | 0.033 | 0.045 |
| recessive | binary | 0.30 | 0.044 | 0.043 | 0.049 | 0.043 | 0.040 | 0.042 |
| recessive | continuous | 0.05 | 0.049 | 0.051 | 0.050 | 0.053 | 0.049 | 0.052 |
| recessive | continuous | 0.10 | 0.048 | 0.052 | 0.057 | 0.051 | 0.044 | 0.053 |
| recessive | continuous | 0.20 | 0.043 | 0.050 | 0.046 | 0.048 | 0.041 | 0.046 |
| recessive | continuous | 0.30 | 0.055 | 0.049 | 0.051 | 0.058 | 0.048 | 0.051 |

### 3.4 Power analysis

We first compared the two p-value aggregation methods, GETAP-ACAT and GETAP-HMP, across various simulation scenarios. The results reveal a consistent pattern: GETAP-ACAT produced equivalent or slightly superior power compared to GETAP-HMP across the choice of various true genetic models (Figures S1 - S6, S17 - S22). The power gain of ACAT is more noticeable (1-2%) in scenarios when the MAF is low, and the true underlying genetic model is additive or dominant (Figures S1, S2, S4, S5, S17, S18, S20, S21). Therefore, given the marginally better performance of GETAP-ACAT compared to GETAP- HMP, we proceed by using the Cauchy p-value combination as the representative strategy for the GETAP approach in subsequent comparisons.

We now summarize the power comparisons stratified by the true underlying genetic model. For each scenario, we compare how GETAP performs relative to: (1) misspecified 1df tests that employ incorrect genotype coding, and (2) the 2df genotypic, model-free test. This framework directly addresses the paper’s central objective: that aggregating evidence across multiple models via GETAP provides robust power.

#### True genetic model is additive

An additive model is the most commonly assumed model. When the true mode of inheritance is additive, the additive GxE test appeared as the most powerful test across various simulation settings for continuous phenotypes, as expected (Figures S7 and S9). We observe a similar pattern for binary phenotypes (Figures S23 and S25). Compared to the true additive model, the misspecified dominant model resulted in a marginal power loss for small MAFs. However, the power loss was greater for larger MAFs, ranging from 1% to 8% for continuous phenotypes (Figures S7 and S9) and 1% to 6% for binary phenotypes (Figures S23 and S25). The misspecified recessive model suffered from maximum power loss, which was most severe at lower MAFs, ranging from 5% to 70% for continuous phenotypes (Figures S7 and S9) and from 5% to 73% for binary phenotypes (Figures S23 and S25). The power loss decreased as the MAF increased, dropping to approximately 1%–25% for larger MAFs.

GETAP incurred only a limited power loss of 1%–5% compared to the correctly specified additive model at small MAFs for both types of phenotypes. But it performed almost identically to the additive model for moderate to large MAFs (Figures S7,S9, S23, and S25). Compared with the dominant model, GETAP has similar power at lower MAFs, but yields a power gain of 1%–6% as the MAF increases (Figures S7, S9, S23, and S25). In comparison with the recessive model, GETAP provides a substantial power gain, particularly at lower MAFs, ranging from 1% to 70% (Figures S7, S9, S23, and S25). Thus, GETAP successfully aggregated the strong signal from the additive model-based test and performed substantially better than the misspecified dominant and recessive models. Compared to the true additive model, it exhibits a limited power loss of 1%–5%. Therefore, when the true model is additive, the proposed GETAP approach performs robustly and competitively with the additive GxE test. GETAP performs substantially better than the other misspecified models. These patterns are consistent across both continuous and binary environmental factors.

#### True model is dominant

When the true mode of inheritance is dominant, the GxE test based on dominant genotype coding is the most powerful for continuous phenotypes, consistently across both continuous and binary environmental variables (Figures S8 and S10). We observe a similar pattern for binary phenotypes (Figures S24 and S26). The commonly used additive model, which is misspecified in this setting, incurred a limited power loss ranging from 1% to 7%, with the loss increasing as the MAF increased (Figures S8, S10, S24, and S26). The recessive model performed poorly, exhibiting substantial power loss relative to the dominant and additive models, ranging from 3% to 75% (Figures S8, S10, S24, and S26). Across all four combinations of phenotype and environmental variable types, the power loss decreased as the MAF increased.

GETAP effectively bridged this gap. It showed a limited power loss compared to the true dominant model for varying MAFs, ranging in 2 – 6%. Its power curve is nearly indistinguishable from that of the misspecified additive model except for small MAFs, where it marginally loses power by 1 – 3% (Figures S8,S10, S24, and S26). Therefore, the GETAP approach performs competitively with the additive model, the second-best performing method in this case. Thus, GETAP performed substantially better than the misspecified recessive model and nearly as well as the additive model. Overall, it is again a robust alternative.

#### True model is recessive

When the true mode of inheritance is recessive, the recessive GxE test is the most powerful for continuous phenotypes, particularly at lower MAFs (0.05 and 0.1), where the homozygous risk genotype is rare but has a strong effect (Figures 2 and S11). This pattern is consistent across both continuous and binary environmental variables. We observe a similar pattern for binary phenotypes, with the recessive GxE test again achieving the highest power under both continuous and binary environmental variables (Figures 3 and S27).

**Figure 2:**
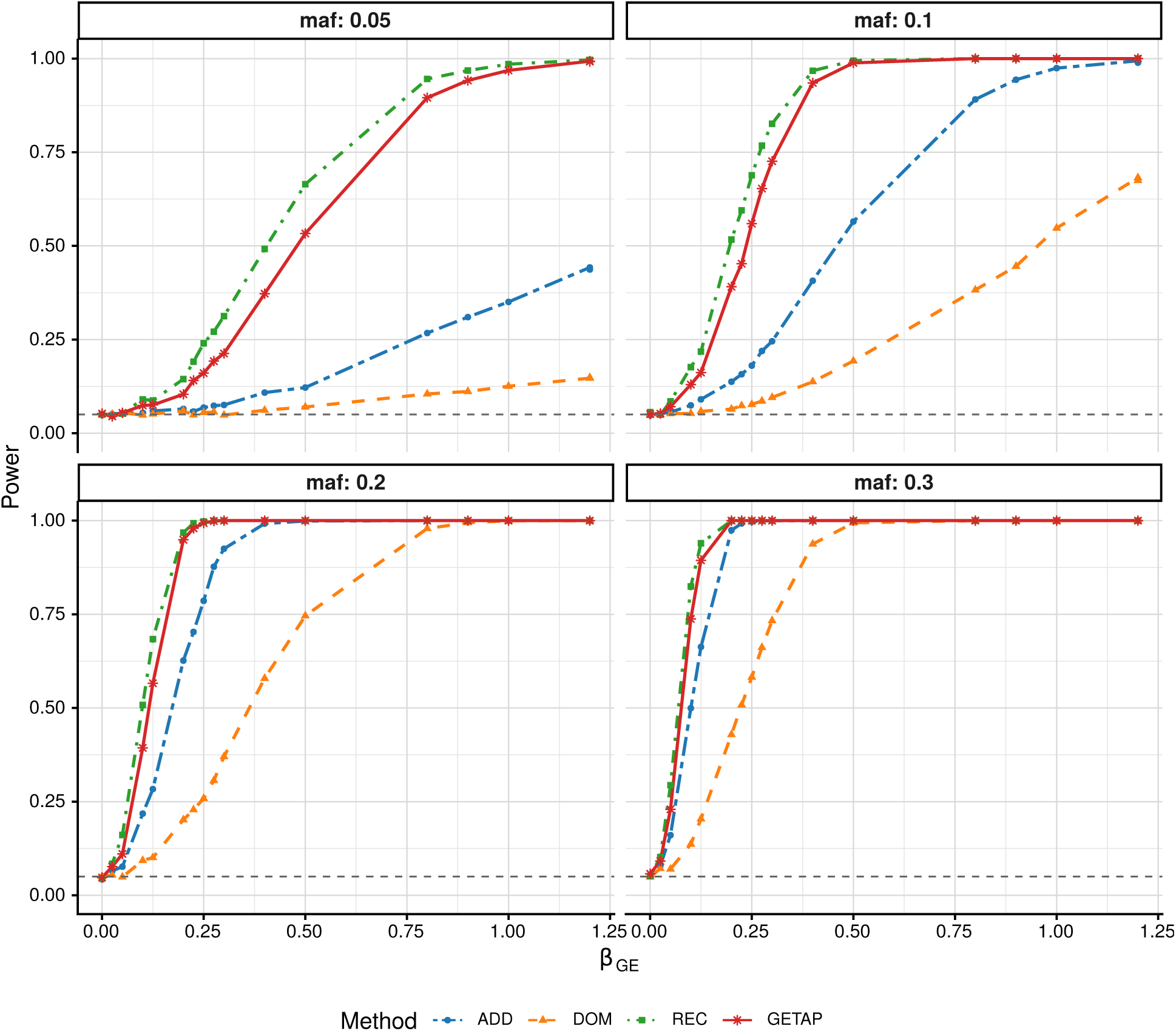
Comparison of statistical power between the 1df tests and GETAP for a continuous phenotype under a recessive genetic model with a continuous environmental exposure. The 1df G×E tests assume an additive (ADD), dominant (DOM), and recessive (REC) genetic model, respectively. Power is empirically estimated at 0.05 level of significance using 2, 000 iterations. The sample comrpises *n* = 10,000 individuals.

**Figure 3:**
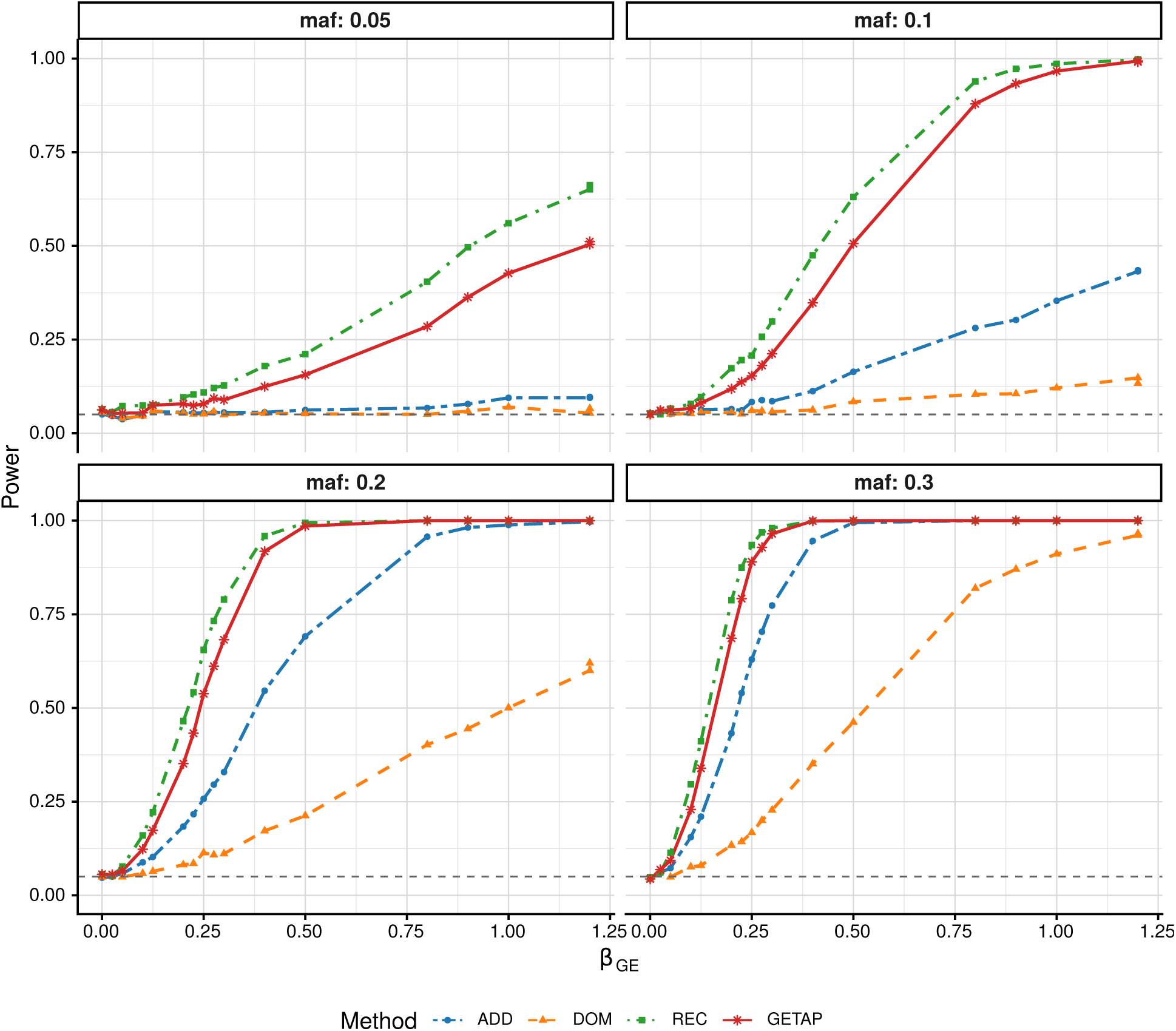
Comparison of statistical power between the 1df tests and GETAP for a binary phenotype under a recessive genetic model with a continuous environmental exposure. The 1df G×E tests assume an additive (ADD), dominant (DOM), and recessive (REC) genetic model, respectively. Power is empirically estimated at 0.05 level of significance using 2, 000 iterations. The sample comrpises *n* = 10,000 individuals.

The additive and dominant models showed severe power loss, as they incorrectly dilute the strong effect present only in the rare homozygous genotypic group. GETAP outperformed these two misspecified alternatives. Specifically, for continuous phenotypes, GETAP achieved a power gain of 2% – 55% over the additive model and 5% – 70% over the dominant model, with the gain decreasing as the MAF increased (Figure 2). The power loss was comparably lower when the environmental variable is binary (Figure S11). We observed a similar pattern for binary phenotypes, where GETAP yielded power gains of 1% – 45% compared with the additive model and 1% – 50% compared with the dominant model (Figures 3, and S27), while other trends remained consistent with those observed for continuous phenotypes. When compared with the true recessive model, GETAP incurred a limited power loss of 1%–13% at small MAFs (Figures 2, and 3); however, this loss diminished for larger MAFs and stronger genetic effects. When the environmental variable is binary, this power loss is comparatively smaller (Figures S11, and S27), although the overall pattern remained similar for both continuous and binary phenotypes.

Therefore, GETAP outperforms the misspecified additive and dominant models in these settings where the true genetic model is recessive. In this scenario, the standard additive GxE test and the dominant test suffer substantial power loss due to model misspecification. By aggregating evidence across different inheritance models, GETAP substantially improves power against misspecified alternatives. However, it incurs a limited power loss relative to the correctly specified recessive model, particularly at low MAFs, where signal detection is most challenging.

#### ACAT versus model-Free 2df Test

We find that when the true genetic model is additive or dominant, GETAP is uniformly more powerful than the 2df test, with power gains ranging from 1%–6% under the additive model and 1%–4% under the dominant model. We observe this pattern for both continuous and binary phenotypes, and across various choices of MAFs and types of environmental factors (Figures 4, S12, S14, S15, S28, S29, S31, S32).

**Figure 4:**
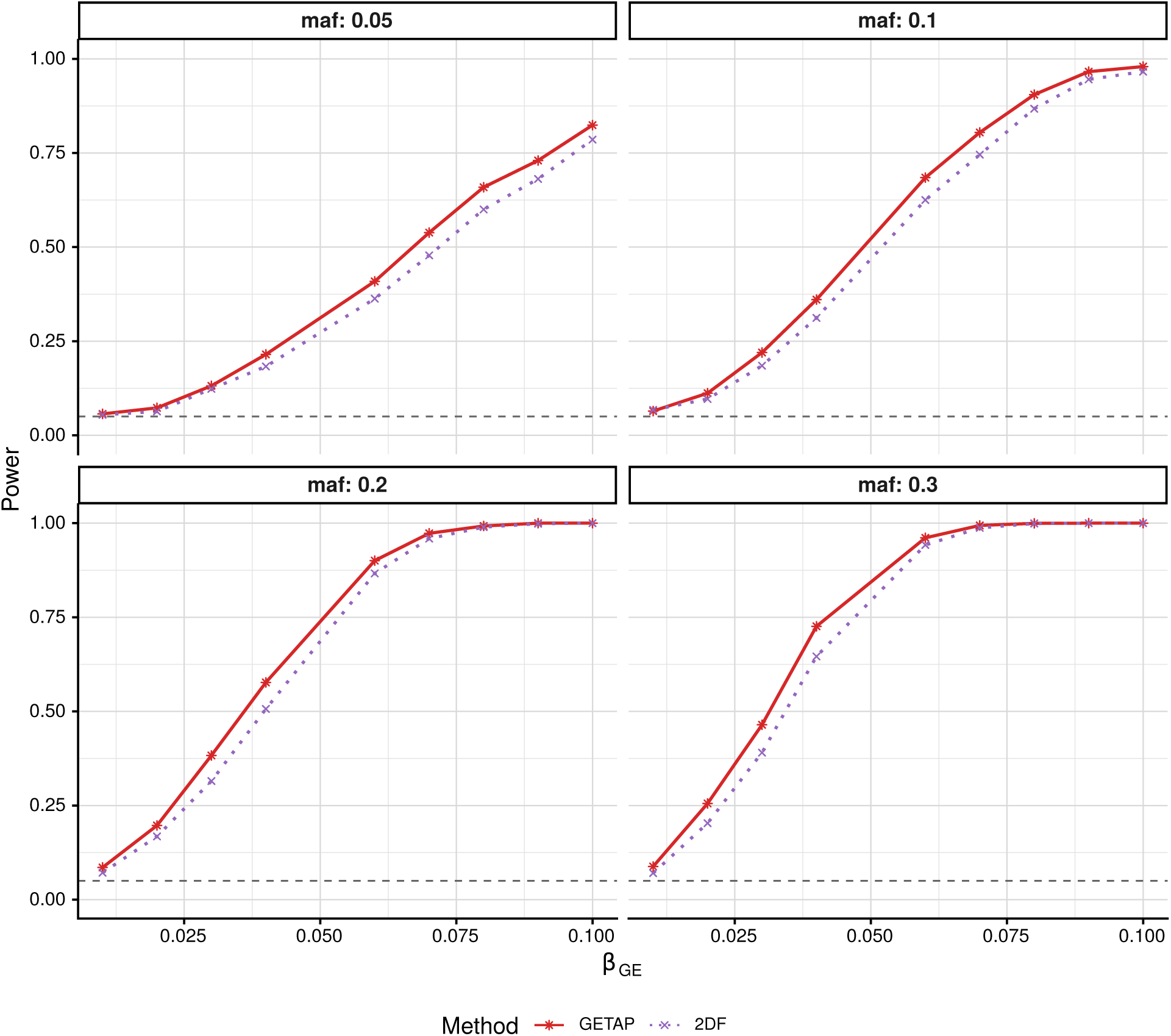
Comparison of statistical power between GETAP and the 2df genotypic GxE test for a continuous phenotype under an additive genetic model with a continuous environmental exposure. Power is empirically estimated at 0.05 level of significance using 2, 000 iterations. The sample comrpises *n* = 10,000 individuals.

When the true model is recessive, the 2df test exhibits a slight power advantage over GETAP at lower MAFs (e.g., 0.05 and 0.1), with gains of approximately 1%–2%. However, this difference diminishes as the MAF increases, and the two methods perform similarly at moderate to larger MAFs, such as 0.2 and 0.3 (Figures S13, S16, S30, S33). Overall, these results demonstrate that GETAP provides a consistently powerful alternative to the 2df test for dominant and additive genetic models, while being competitive in recessive settings as well.

In summary, the simulation results confirm that the GETAP approach via ACAT p-value combination offers good power robustly across various true genetic models for both continuous and binary phenotypes. It offers substantially higher power than the commonly used additive model when the true genetic model is recessive. When the true model is additive or dominant, GETAP incurs a marginal power loss relative to the additive model in some scenarios, particularly for lower MAFs. The method offers higher power than the 2df test when the true genetic model is additive or dominant; for a recessive model, it performs similarly or marginally loses power. Overall, GETAP is robustly powerful for detecting GxE interactions while controlling the type I error rate adequately.

## 4 Real data application

We evaluated the proposed GETAP approach using large-scale data from the UK Biobank (UKB), a population-based cohort with extensive genetic, phenotypic, and environmental characterization. The UKB provides genome-wide genotype data for approximately 500, 000 participants, together with harmonized measurements of numerous quantitative and disease-related phenotypes and a wide range of lifestyle and environmental factors. This excellent resource enables us to directly assess whether the robustness and power gains of the GETAP approach observed in simulation studies extend to GxE analyses in a realworld setting, where genetic architectures of complex phenotypes and diseases are heterogeneous, and environmental exposures are measured with varying precision.

We analyzed several different phenotype-environment combinations spanning both continuous and binary outcomes. Continuous phenotypes include biomarkers of glycemic control, pulmonary function, adiposity, and metabolic and inflammatory processes, paired with smoking-related and dietary environmental exposures. We also considered behavioral or lifestyle factors. Binary outcomes include type 2 diabetes and chronic obstructive pulmonary disease (COPD), analyzed in conjunction with sleep duration and cumulative smoking exposure, respectively. We provide detailed definitions of all phenotypes, environmental exposures, and inclusion criteria in the Supplementary Information.

For each phenotype-environment pair, we curated high-quality datasets using strict quality control procedures, focusing on White British ancestry with complete covariate information. We performed the analyses on the UK Biobank Research Analysis Platform (RAP), a cloud computing environment. We used autosomal genotype calls. We excluded SNPs with minor allele frequency below 1% since sample size is very large, followed by Hardy–Weinberg equilibrium (HWE) screening (HWE test *P <* 5 × 10*^-^*^6^), and exclusion of SNPs with genotyping call rate below 95%. After implementing the genotype QC steps, we obtained a common set of approximately 6, 05, 000 autosomal SNPs for genome-wide analysis of each phenotype-environment combination. We adjusted the phenotype for relevant covariates: age, age^2^, sex, and the top ten genetic principal components (PCs), with additional adjustment for medication use wherever relevant and applicable. We applied the standard rank-normal transformation to continuous outcomes to achieve normality. We defined binary disease phenotypes using validated algorithms in UKB, integrating hospital records, primary care data, medication information, and self-reported diagnoses.

Across all analyses, we performed genome-wide tests of GxE interaction using additive, dominant, and recessive genotype encodings, as well as the model-free 2-degree-of-freedom(2df) genotypic test. We compared these results with the proposed GETAP approach, which aggregates evidence across different genotype encodings using the Cauchy combination of p-values to provide a genetic-model-robust test of interaction. Since GxE signals are limited due to small interaction effect sizes, we applied the Benjamini-Hochberg (BH) FDR control procedure with FDR *<* 0.05 to identify genome-wide (GW) significant signals. Next, to identify independent interaction signals, we applied LD-based clumping to the marginally significant variants using PLINK v1.9, with a 500 kb window and an LD threshold of *r*^2^ *<* 0.2. We performed LD clumping separately for each method, enabling direct comparison of independent loci identified by the different tests. We provide the quantile-quantile (QQ) plots to assess calibration and genomic inflation.

In the following subsections, we discuss the results on GxE interactions for different phenotype environmental factor combinations. We first describe the results for continuous phenotypes, followed by binary disease phenotypes.

### 4.1 Gene-environment interactions underlying Glycated hemoglobin (HbA1c)

Glycated hemoglobin (HbA1c; UKB Data-Field 30750) reflects long-term blood glucose control and is a biomarker of type 1 and 2 diabetes. Measurements were obtained at baseline using high-performance liquid chromatography and reported in mmol/mol units. We excluded individuals with implausible HbA1c levels (*<* 15 or *>* 200 mmol/mol), type 1 diabetes, pregnancy, or conditions affecting erythrocyte turnover. The final sample comprised 3, 00, 599 unrelated individuals of White British ancestry.

#### 4.1.1 HbA1c and Cumulative Smoking Exposure (Pack-Years)

Cumulative smoking exposure was quantified using lifetime pack years (PackSMOKE; UKB Data-Field 20161), computed from cigarettes smoked per day and smoking duration following UK Biobank procedures. A numeric value for pack-years was assigned to all ever-smokers and set to zero for never-smokers, yielding a continuous exposure defined across all participants in the full cohort.

Testing for genome-wide GxE interactions led to substantial differences in the number of GxE signals detected by the five methods. The additive, dominant, and recessive 1df tests identified 25, 10, and 58 significant SNPs with GxE interaction, respectively, while the 2df genotypic test detected 45 interactions. The GETAP approach identified 87 significant SNPs with GxE interaction, substantially more than the other approaches (Figure 6). A five-way Venn diagram (Figure 5) shows the varying extent of overlap in the number of discoveries among the five approaches. For example, the GETAP approach identified a common subset of 25, 10, 39, and 38 SNPs with the 1df additive, dominant, recessive tests and 2df test, respectively (Figure 5). We note that the GETAP approach identified 20 SNPs with GxE interaction, which were not detected by any single-model or 2df test. This highlights the advantage of p-value aggregation in recovering GxE interaction signals that individual genetic models may not capture adequately. For example, SNP rs1045997 on chromosome 19 shows consistent evidence of GxE interaction in the additive and dominant models (*P* = 2.3 × 10*^-^*^6^ and 1.3 × 10*^-^*^6^); but these do not survive the multiple-testing correction in any single-model or 2df test. However, GETAP identifies this SNP as a significant GxE SNP by aggregating these partial signals. The 2df test can miss such signals primarily because it spends an extra degree of freedom.

**Figure 5:**
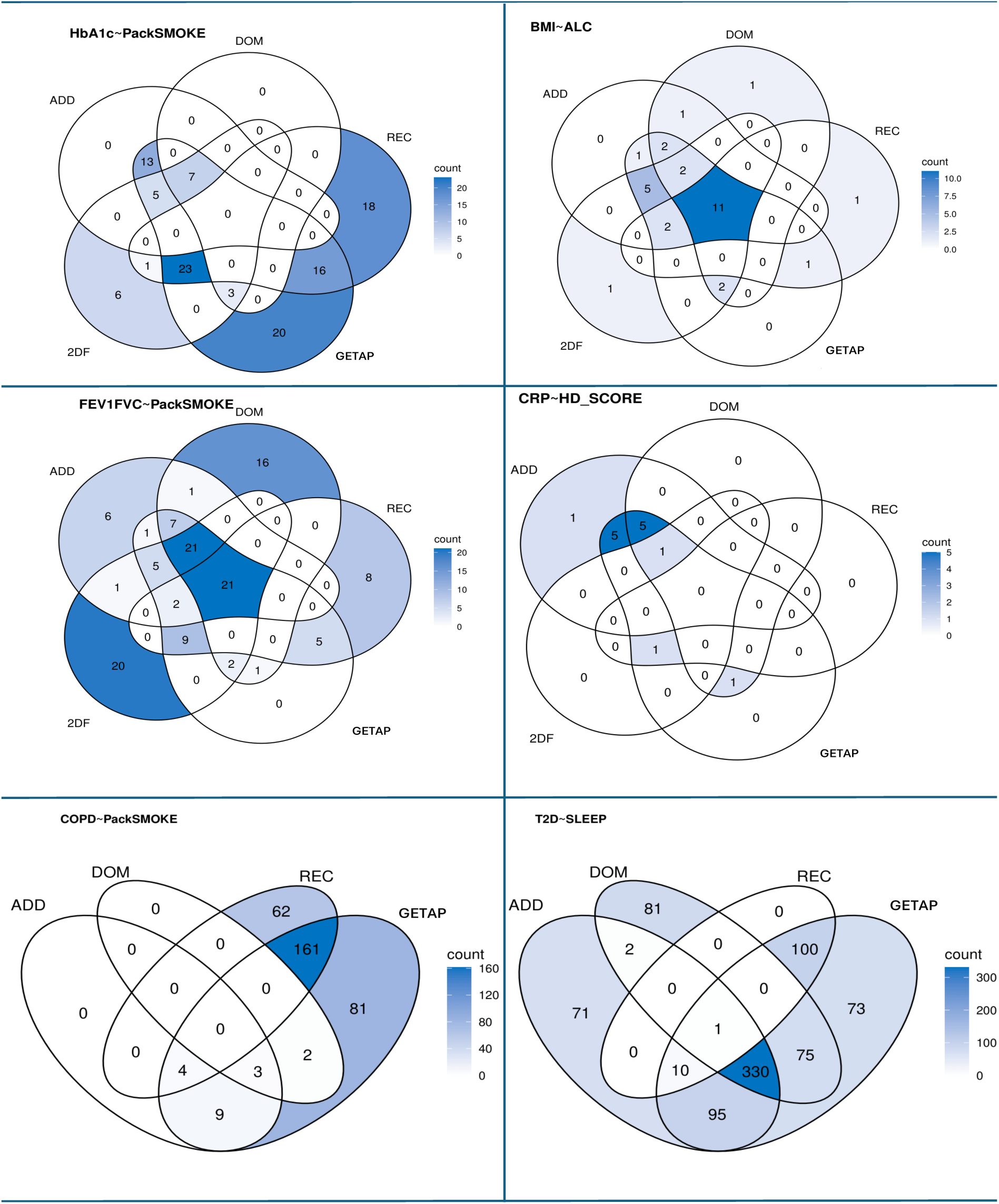
Venn diagrams showing overlap of significant GxE SNPs (FDR *<* 0.05) identified by additive (ADD), dominant (DOM), recessive (REC), 2-degree-of-freedom (2DF) genotypic, and the proposed GETAP tests for multiple phenotype–environment pairs. The numbers in the intersection regions represent the number of SNPs detected by multiple approaches. Degree of shading intensity corresponds to SNP counts.

**Figure 6:**
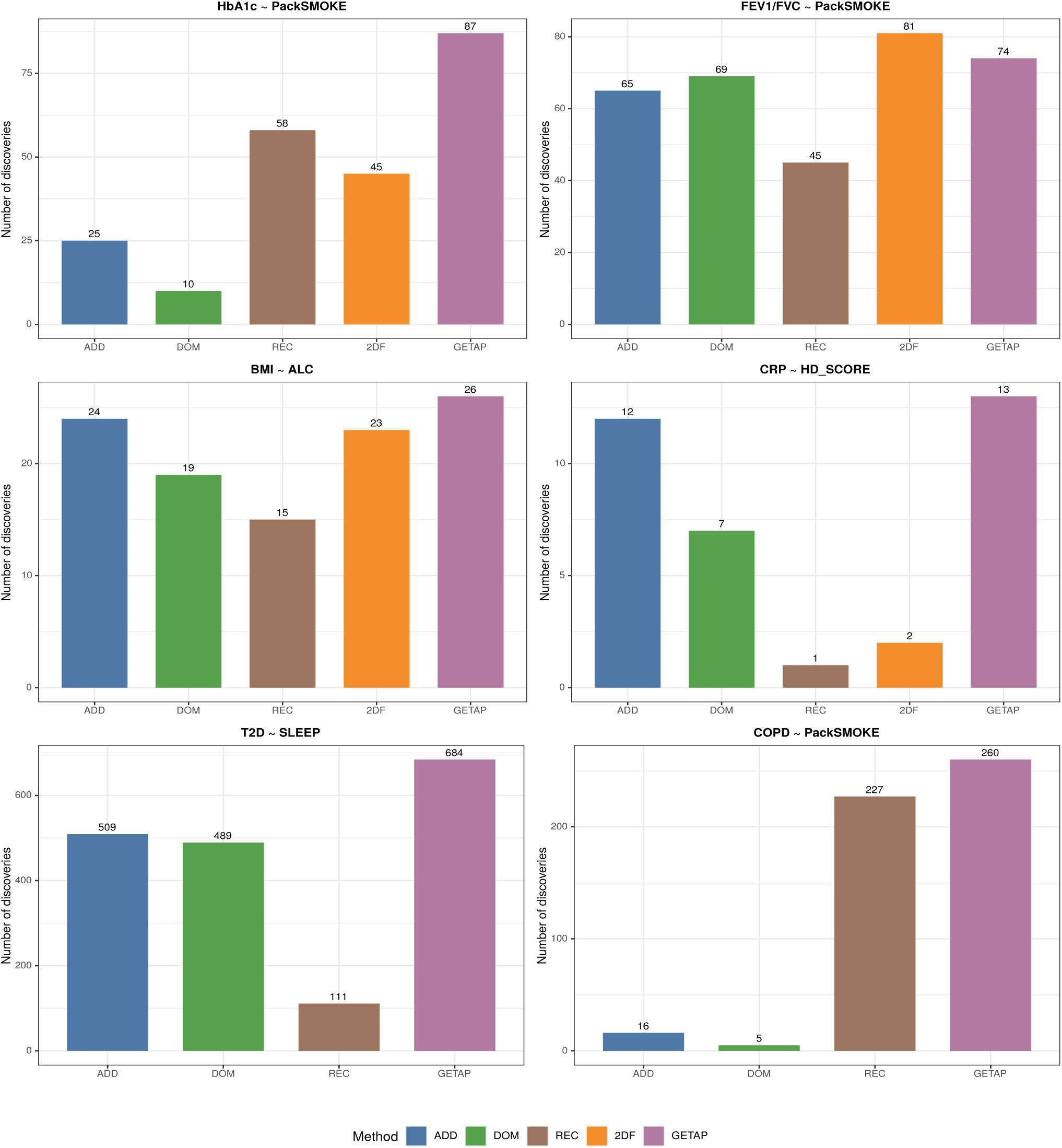
Bar plots showing the number of significant G×E interaction SNPs identified at 5% FDR for six different selected phenotype-environment pairs (HbA1c ∼ PackSMOKE, FEV1/FVC ∼ PackSMOKE, BMI ∼ ALC, CRP ∼ HD SCORE, T2D ∼ SLEEP, and COPD ∼ PackSMOKE). Bars represent discovery counts for the additive (ADD), dominant (DOM), recessive (REC) 1df tests, two-degree-of-freedom genotypic test (2DF), and the proposed GETAP approach.

LD-based clumping identified 24 independent GxE loci from the 25 SNPs marginally identified by the additive model, 8 independent GxE loci from 10 SNPs marginally identified by the dominant model, 55 loci from 58 recessive signals, 42 loci from 45 2df signals, and 82 loci from the 87 GETAP-identified signals. Thus, the majority of GxE interaction signals mapped to weakly correlated genomic regions, suggesting limited redundancy due to LD. We note that the LD tagging pattern of GxE interaction effects can be distinct from that of main genetic effects. The GETAP approach yielded the largest number of independent GxE loci, with the strongest signal at rs407423 on chromosome 8 (*P* = 1.54 × 10*^-^*^9^). The additive model identified 24 loci led by rs62395369 on chromosome 6 (*P* = 1.58 × 10*^-^*^7^), while the dominant model detected 8 loci, led by rs10062026 on chromosome 5 (*P* = 5.71 × 10*^-^*^8^). The recessive and 2df analyses also highlighted rs407423 on chromosome 8 as the top GxE signal. The latter demonstrates the contribution of a non-additive genetic model in detecting G×smoking interactions underlying the HbA1c level. We describe these independent GxE loci in the supplementary data files.

We evaluated the genomic inflation factor (GIF or *λ*) across the different tests to assess possible inflation in the test statistics. We found the GIF to be 1.29 for the recessive model, and 1.31 for the additive and dominant models. We obtained a higher inflation for the 2df test: *λ* = 1.45 (Figure S34). In comparison, GETAP yielded a lower GIF: *λ* = 1.27. We note that the GIF is more than one, primarily due to polygenic interaction effects present for this phenotype-environment combination. We have observed estimates of GIF around one for other pairs of phenotype-environmental factors for which all methods identified substantially fewer GxE interaction signals (Table S4).

### 4.2 GxE interactions for Pulmonary Function related phenotypes

The UK Biobank assessed pulmonary function using the ratio of forced expiratory volume in one second (FEV1) to forced vital capacity (FVC): FEV1/FVC. Spirometry-derived FEV1 (Data-Field 3063) and FVC (Data-Field 3062) were measured at baseline following standardized ATS/ERS protocols. We restricted the analyses to high-quality spirometry measurements and excluded participants with conditions known to affect lung function severely. We standardized measurements using Global Lung Initiative reference equations and removed extreme outliers (|*z*| *>* 5).

#### 4.2.1 FEV_1_/FVC and Cumulative Smoking Exposure (Pack-Years)

We examined gene-environment interactions between pulmonary function and cumulative smoking exposure by analyzing the FEV_1_/FVC ratio and lifetime pack years (PackSMOKE) combination. Here, we regard PackSMOKE as the lifestyle factor. The final analytic sample comprised 294, 405 unrelated individuals of White British ancestry. At 5% FDR, all five methods identified a substantial number of significant interaction signals. The additive, dominant, and recessive tests detected 65, 69, and 45 SNPs, respectively. The 2df genotypic test identified 81 SNPs with GxE interaction. The GETAP-ACAT procedure identified 74 significant SNPs (Figure 6). GETAP identified 17 signals not detected by the additive model and 22 not detected by the dominant model, while maintaining substantial overlap with all single-model tests, indicating effective aggregation across heterogeneous genotype encodings.

The overlap structure across methods, summarized in a five-way Venn diagram (Figure 5), reflects a complex overlapping pattern. Notably, each of the three 1df tests and the 2df test identified some unique SNPs which were missed by the others. However, the GETAP approach did not identify any unique SNPs (Figure 5). But GETAP detected more SNPs than the 1df tests. The 2df test identified a few more SNPs than GETAP. Overall, GETAP performed competitively for this combination. We note that the 2df test detected more unique SNPs here. This indicates that the underlying genetic model can sometimes be uncommon, not necessarily one of additive, dominant, or recessive.

Evaluation of genomic inflation showed moderate inflation across all methods (Figure S36), which we attribute to polygenic GxE effects for this phenotype-environmental factor combination. Estimated GIF is similar for the 1df additive and dominant tests: *λ* = 1.25, and slightly lower for the recessive test: *λ* = 1.22. In comparison, GETAP exhibited a lower inflation factor: *λ* = 1.19 (Table S4). The 2df test yielded the highest GIF: *λ* = 1.36, mainly due to being more powerful for this pair.

LD-based clumping identified 40 independent GxE loci from 74 GETAP-significant variants found by the BH FDR rule with *P <* 5.95 × 10*^-^*^6^. On chromosome 15, rs16969968 had the strongest signal with *P* = 1.81 × 10*^-^*^17^. This SNP resides within the *CHRNA* gene cluster. We identified 30 independent GxE loci from 65 SNPs detected by the additive test. Similarly, we identified 11 GxE loci from 29 variants under the dominant model, and 23 GxE loci from 45 variants under the recessive model. The 2df test yielded 48 independent GxE loci from 81 variants, more than the others. Together, these results demonstrate that many genetic variants interact with cumulative smoking exposure to impact the pulmonary function interactions. Our proposed GETAP approach performed well for this combination of phenotype and exposure. We provide the independent GxE loci in the supplementary data.

### 4.3 GxE interactions underlying adiposity-related phenotype

In the UK Biobank, body mass index (Data-Field 21001) was calculated from height and weight measurements obtained for the participants during their initial assessment centre visit. We consider body mass index (BMI) as the phenotype.

#### 4.3.1 BMI and alcohol intake frequency

We regard alcohol intake frequency as the environmental exposure for studying gene-environment interactions for BMI. Alcohol intake frequency (Data-Field 1558) was assessed via the touchscreen questionnaire and modeled as an ordinal exposure ranging from “never” to “daily or almost daily”. We include the details of the conversion of the alcohol intake frequency to numeric values in the supplementary material. After quality control, the BMI and alcohol intake frequency analysis included 379, 836 participants.

At 5% FDR, all five methods identified a moderate number of significant GxE interactions. The additive, dominant, and recessive 1df tests detected 24, 19, and 15 significant SNPs, respectively. The 2df genotypic test identified 23 SNPs with significant interaction effects. The GETAP approach yielded 26 significant SNPs, slightly more than the others (Figure 6). GETAP recovered some SNPs missed by the additive, dominant, and recessive models, indicating improved sensitivity to detect interaction effects that a specific genotype encoding may not capture well. Overlapping patterns across methods (Figure 5) revealed that multiple different tests consistently identified a common set of 11 SNPs. GETAP aggregated signals arising from all three different genetic modes of inheritance, rather than favoring any single model. Minimum *p*-values among the FDR-significant SNPs were in the order of 10*^-^*^11^ for the additive, GETAP, and 2df tests, which indicates some strong interaction signals.

LD-based clumping identified 10 independent loci from 25 GETAP-significant variants. The strongest signal was *rs*17817449 on chromosome 16 (*P* = 3.04 × 10*^-^*^11^) near the *FTO* gene, with 16 tagged variants. The FTO gene is well-known to regulate BMI. The additive model-based analysis also identified 10 GxE loci from 24 variants, led by the same SNP (*P* = 1.02 × 10*^-^*^11^). The dominant model detected 8 independent loci from 19 variants, with *rs*1421085 on chromosome 16 as the top signal (*P* = 1.43 × 10*^-^*^7^), while the recessive model identified 3 loci from 15 variants. The 2df test detected 8 independent loci from 23 variants. We provide the independent GxE loci detected by our approach, GETAP, in the supplementary data.

Evaluation of genomic inflation showed a limited inflation across all methods (Figure S38). Estimated GIF for the additive, dominant, and recessive tests are *λ* = 1.16, 1.15, 1.14, respectively. The GIF is marginally higher for the 2df test: *λ* = 1.2. In comparison, GETAP exhibited lower inflation: *λ* = 1.12, suggesting an improved calibration while maintaining competitive power.

### 4.4 Gene-environment interactions for Metabolic and Inflammatory Biomarkers

We also analyzed inflammatory and metabolic biomarkers: C-reactive protein (CRP; Data-Field 30710) and triglycerides (TG; Data-Field 30870), measured using enzymatic assays at baseline. In the TG analysis, we adjusted for lipid-lowering medication. CRP and TG were each paired with the Healthy Diet Score (HD-SCORE). HD-SCORE is a composite metric summarizing four dietary components: fruit and vegetable intake, fish consumption, red meat intake, and processed meat intake, constructed following American Heart Association guidelines. Each component was dichotomized according to recommended thresholds, yielding a score ranging from 0 to 4, with higher values indicating healthier dietary patterns. Final analytic sample sizes for CRP and TG analyses were approximately 362, 698 and 363, 187 individuals of White British ancestry, respectively.

#### 4.4.1 C-Reactive Protein and Healthy Diet Score

We investigated gene-environment interactions considering serum C-reactive protein (CRP) levels as the phenotype and the dietary quality using the Healthy Diet Score (HD-Score) as the lifestyle factor. At 5% FDR, the additive, dominant, and recessive 1df tests identified 12, 7, and 1 significant SNPs, respectively. The 2df genotypic test detected 2 signals. The GETAP procedure identified 13 significant SNPs, more than the others. Overlap patterns across methods (Figure 5) revealed both shared and method-specific signals. GETAP again produced more signals than the other approaches (Figure 6).

Assessment of genomic inflation across all methods (Figure S39) exhibited adequate covariate adjustment. Genomic Inflation factors are around unity for the additive, dominant, and recessive 1df tests: *λ* = 1.01, 1.01, 0.98. We observed limited deflation for the 2df test and GETAP: *λ* = 0.94, 0.9, respectively. We note that the number of GxE signals obtained for this phenotype-environmental exposure is lower than the above combinations. Proportionately, we also observe a lower GIF for the methods for the current combination (Table S4). This indicates a more polygenic interactions for the above pairs of phenotype-environment.

LD-based clumping identified 3 independent loci from 13 GETAP-significant variants (*P <* 9.7 × 10*^-^*^7^). The lead signals were rs876537 on chromosome 1 near the *CRP* gene (*P* = 5.08 × 10*^-^*^9^), rs12145237 on chromosome 1 (*P* = 3.58 × 10*^-^*^8^), and rs79709269 on chromosome 15 (*P* = 5.14 × 10*^↑^*^8^). The additive and dominant tests identified 2 independent loci each. The recessive and 2df tests detected one and two loci, respectively. Overall, the CRP-diet analysis represents a weak GxE signal setting. However, GETAP again provided a modest but consistently higher number of GxE signals. We have included the GxE loci in the supplementary data.

Even though we have discussed the results for the above combinations, which have a moderate number of GxE signals, we did not obtain any signal for many other combinations of phenotype-environment (Table S5). For these combinations, we obtained a genomic inflation factor of around 1, which again justifies the adequacy of the covariate adjustment in the genome-wide GxE studies that we have performed. We performed the GxE analysis for some other phenotype-environment combinations. For brevity, we have discussed them in the supplementary information.

### 4.5 Analysis of binary phenotypes

Type 2 diabetes (T2D) is a complex metabolic disorder with a well-characterized genetic architecture and strong associations with lifestyle-related factors, including sleep patterns. Another important binary phenotype is chronic obstructive pulmonary disease (COPD), a respiratory phenotype with a strong environmental component, e.g., smoking. Therefore, we analyze T2D and COPD for detecting GxE interactions while considering sleep duration and cumulative smoking exposure as the lifestyle factors, respectively.

#### 4.5.1 T2D and Sleep duration

UK Biobank defined T2D case-control status using a validated multimodal algorithm that integrates hospital inpatient records, primary care data, medication usage, and self-reported history. We identified cases based on ICD-10 diagnostic codes (E11, E13, E14), self-reported diabetes excluding type 1 diabetes, and the use of glucose-lowering medications (e.g., metformin, sulfonylureas, SGLT2 inhibitors). We excluded individuals with type 1 diabetes (E10), gestational diabetes, or early insulin initiation indicative of autoimmune diabetes. We defined participants without diabetes-related diagnosis codes, no diabetes medication use, and no self-reported diabetes as controls. After curating cases and controls, ancestry restriction to unrelated White British individuals, and standard sample quality control, the final dataset comprised 20, 726 T2D cases and 275, 757 controls. We note that the possible over-representation of controls in the sample will not decrease power.

We incorporated sleep duration as the lifestyle factor, derived from baseline self-reported UKB data (Field ID 1160), reflecting the average number of hours slept in a 24-hour period, including naps. We treat sleep duration as a continuous environmental variable and exclude individuals reporting implausible average sleep durations (*<* 2 or *>* 17 hours per day) or who have missing data. Given the binary nature of the phenotype, we performed logistic regression to evaluate GxE interactions using 1df tests under additive, dominant, and recessive genotype coding using PLINK. We combined the GxE p-values obtained from the three different tests of interaction using the Cauchy aggregation to obtain a single p-value for each SNP. We could not include the 2-degree-of-freedom genotypic test for binary phenotype analyses due to our limited resources to perform genetic analyses in the UK Biobank research analysis platform (cloud computing).

At 5% FDR, the additive, dominant, and recessive tests identified 509, 489, and 111 significant SNPs to have a GxE interaction with sleep duration, respectively. The GETAP approach detected 684 significant SNPs, 175 SNPs more than the additive model, which is the best-performing 1df test here (Figure 6). Among 684 GETAP-significant SNPs, 436 overlapped with those found by the 1df additive test, 406 overlapped with those detected by the dominant test, and 111 overlapped with the recessive-model signals (Figure 5). Hence, GETAP captured a major proportion of signals identified by the additive and dominant models, and all detected by the recessive model. Importantly, GETAP uniquely identified 73 SNPs which the 1df tests missed. GETAP identified many of these SNPs by aggregating limited evidence provided by the individual 1df tests, which could not pass the multiple testing correction.

To characterize independent GxE loci while reducing redundancy due to LD, we performed LD-clumping on the sets of SNPs passing FDR correction for each method. The 509 additive-model SNPs collapsed into 414 clumps, hence GxE loci. The dominant-model SNPs yielded 419 loci from 489 SNPs. The recessive-model signals resulted in 95 GxE loci from 111 SNPs. The GETAP-detected 684 SNPs led to 563 independent GxE loci. Several GETAP-identified LD clumps did not overlap with those detected by any 1df single genetic model-based test. Overall, GETAP performed substantially better than the 1df single genetic model-based tests for the type 2 diabetes and sleep duration pair.

We next evaluate the genomic inflation using quantile-quantile (QQ) plots and genomic inflation factor (GIF) (Figure S41). We obtained the estimated GIFs as *λ* = 1.48 for the additive model-based test, *λ* = 1.48 for the dominant model-based test, *λ* = 1.43 for the recessive test, and *λ* = 1.59 for GETAP. This is consistent with the pattern of finding higher GIFs when the number of GxE signals is higher, reflecting more polygenic GxE architecture (Table S4). We further explore the sample-size-standardized genomic inflation factor because the current case-control dataset is large, which can also influence the increase in GIF. This phenomenon is expected to be more common in a case-control genetic study [13], even under well-calibrated polygenic architectures. We report the sample-size-standardized GIF, *λ*_1000_, corresponding to an equivalent study of 1, 000 cases and 1, 000 controls [13]. Following the procedure, we obtained *λ*_1000_ rescaling the observed inflation factor, *λ*_obs_, using the effective sample size:

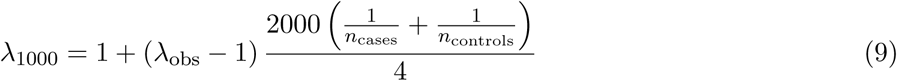

For the current analysis of T2D-sleep duration, we obtained *ϱ*_1000_ as 1.02. Since the sample size standardized GIF is close to unity, we discard any possible role of residual population stratification, cryptic relatedness, or other unadjusted covariates in the observed inflation. We rather attribute the inflation of the primary GIFs to the polygenic GxE architecture of T2D.

#### 4.5.2 COPD and Pack-years of smoking

We next analyzed COPD, taking pack-years of smoking as the lifestyle factor or environmental exposure. The UK Biobank defined the COPD cases and controls using a combination of hospital inpatient records, self-reported diagnoses, and spirometry-based exclusion criteria. After phenotype definition, ancestry restriction to unrelated White British individuals, and standard quality control procedures, the final analysis included 23, 033 COPD cases and 2, 49, 599 controls. Again, a possible over-representation of controls in the sample should not decrease power.

We conducted genome-wide GxE interaction analyses using logistic regression. We implemented the same steps as performed for the T2D analysis above. The additive and dominant 1df tests identified 16 and 5 significant SNPs with GxE interaction, respectively. The recessive test identified 227 significant SNPs with GxE interaction, which is substantially larger than the additive and dominant models. The GETAP approach detected 260 significant SNPs with GxE signal, offering more signals than the recessive model-based test, the best performing 1df test here (Figure 6). The overlap pattern indicates that GETAP captured the majority of GxE signals identified by 1df tests (Figure 5). In addition, GETAP uniquely detected 81 SNPs that others missed.

LD clumping produced 13 GxE loci from 16 marginally significant SNPs obtained from the additive model, 5 GxE loci from 5 dominant model SNPs, 200 GxE loci from 227 recessive model SNPs, and 219 GxE loci from 260 GETAP identified SNPs. We obtained the estimated genomic inflation factors as *ϱ*_obs_ = 1.3 for the additive model, 1.31 for the dominant model, 1.2 for the recessive model, and 1.27 for GETAP (Figure S42). After sample size standardization, the corresponding *ϱ*_1000_ values are 1.01 for all the approaches. It demonstrates that the observed inflation is again due to polygenic GxE interactions.

### 4.6 Functional annotation and biological interpretation of GxE loci

We next conducted downstream functional genomic analyses, such as gene prioritization and pathway enrichment analysis using the FUMA web-platform [14]. We performed these analyses for two example phenotypes, one continuous: HbA1c level, and one binary: type 2 diabetes. For HbA1c level, cumulative smoking exposure is the lifestyle factor, and for type 2 diabetes, sleep duration is the environmental factor.

#### 4.6.1 GxE SNPs for HbA1c level and cumulative smoking exposure

LD-clumping (*r*^2^ *<* 0.2, 500 kb window) yielded 81 independent GxE loci distributed across the genome. Functional annotation of the candidate SNPs demonstrates that the majority of the GxE variants are located in the noncoding genomic regions, with major enrichment in intronic and intergenic regions (Figure S47B). This pattern suggests that G×smoking effects on long-term glycemic regulation are primarily mediated through regulatory mechanisms influencing gene expression, rather than through protein-altering coding variation.

We next conducted MAGMA gene-based analysis for the candidate SNPs as implemented in FUMA [14]. This highlighted multiple linked candidate genes. We found these genes relevant to metabolic transport, extracellular matrix organization, and inflammation-related signaling pathways (Figure S47A). For example, GxE loci linked with the *SLC19A1* gene suggest a plausible connection between smoking exposure, cellular transport processes, and glycemic burden. It is consistent with the known role of environmental stressors in modifying metabolic susceptibility.

Regional association plots generated using LocusZoom.js [15] also support these findings by demonstrating localized interaction signal peaks residing within LD blocks at multiple independent loci. Representative signals include an interaction locus near the *SLC19A1* gene on chromosome 21 (Figure S47C), as well as additional loci on chromosome 8 (Figure S47D), and on chromosome 5 near the *ADGRV1* gene (Figure S47E). These highlight possible heterogeneous genetic architecture through which cumulative smoking exposure may amplify or attenuate genetic susceptibility, influencing HbA1c levels. We then evaluated pathway-level enrichment of the genes mapped by FUMA using the Reactome method [14]. No pathways reached statistical significance after FDR correction (FDR-adjusted *P <* 0.05). One possible reason is that the total number of GxE SNPs was lower, hence fewer genes were mapped.

#### 4.6.2 GxE SNPs for T2D and sleep duration

LD-clumping (*r*^2^ *<* 0.2, 500 kb window) resulted in 563 independent GxE loci for T2D and sleep duration identified by GETAP. Again, the GxE variants are located in the noncoding genomic regions, with most enrichment in intronic and intergenic regions (Figure 7C). Thus, G↑sleep interaction effects are primarily mediated through regulatory mechanisms influencing gene expression. Gene prioritization analysis using MAGMA highlighted several prioritized candidate genes, including *GRHL1*, *LAMA2*, and *P2RX7* (Figure 7A). These genes implicate pathways related to transcriptional regulation, extracellular matrix remodeling, and immune-metabolic signaling, suggesting potential mechanisms through which sleep duration may modify genetic susceptibility to T2D.

**Figure 7:**
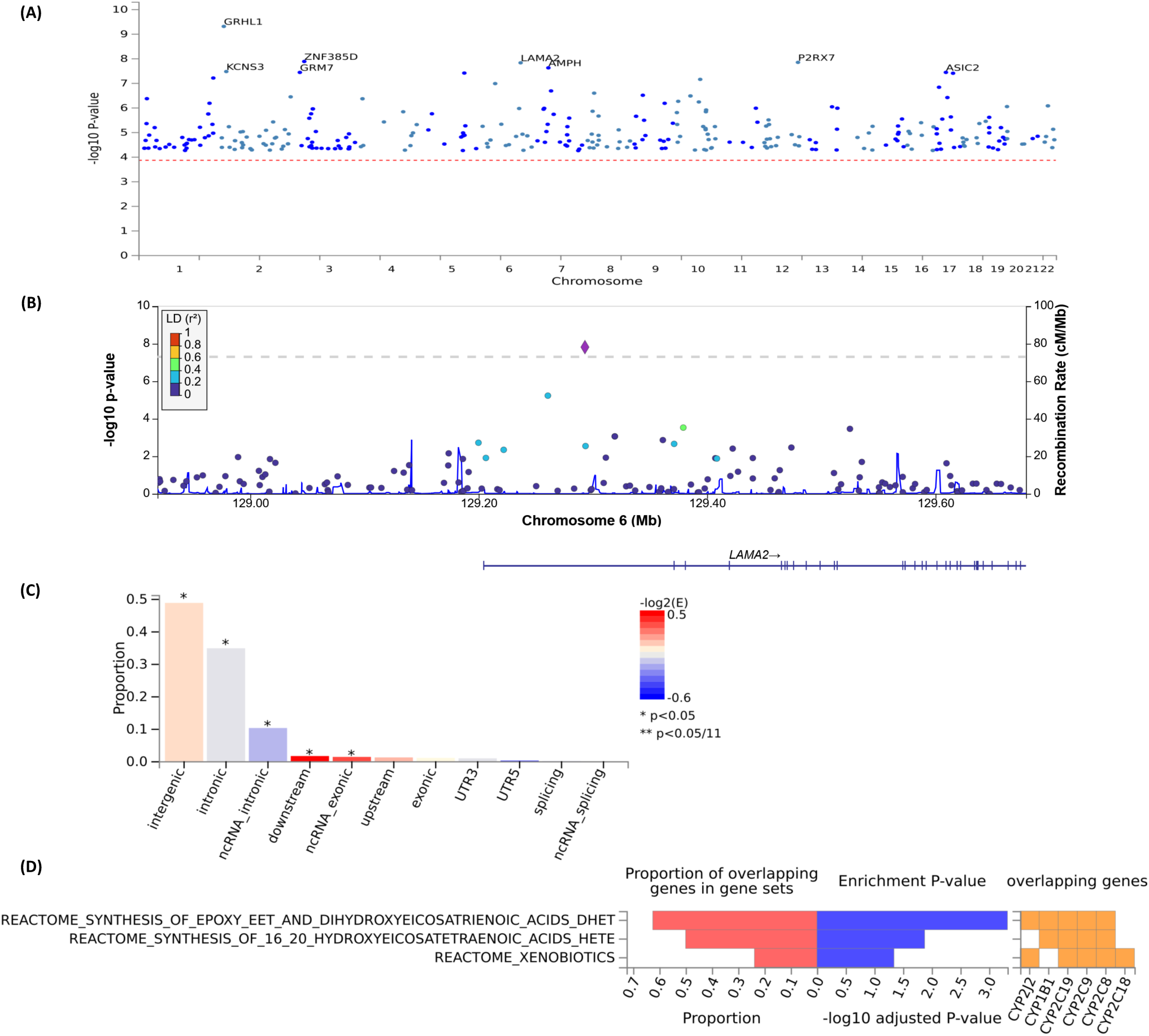
Functional annotation and biological interpretation of gene-environment interaction loci identified by GETAP for type 2 diabetes and sleep duration. (A) Gene-level prioritization by MAGMA derived from GETAP-significant GxE interaction variants, highlighting selected genes across the genome, including *GRHL1*, *LAMA2*, and *P2RX7*. (B) Regional association plot for a representative GxE interaction locus on chromosome 6 near the *LAMA2* gene, showing SNP-level G × E interaction signals, local linkage disequilibrium structure, recombination rate, and genomic context. (C) Functional annotation of mapped GxE variants, showing enrichment in noncoding regions, particularly intergenic and intronic categories, consistent with regulatory mechanisms. (D) Pathway enrichment analysis of genes mapped from GxE loci using Reactome, highlighting significant enrichment for xenobiotic metabolism and lipid-related pathways.

Gene-set enrichment analysis using Reactome highlighted pathways involved in xenobiotic metabolism mediated by cytochrome P450 enzymes, as well as eicosanoid lipid synthesis pathways (Figure 7D). The xenobiotic metabolism pathway reflects processes responsible for detoxification of environmental and endogenous compounds through CYP enzymes, which are known to influence drug metabolism, oxidative stress responses, and metabolic homeostasis [16]. This is particularly relevant in the context of sleep disturbance, which has been linked to altered metabolic clearance, inflammation, and insulin resistance [17]. In addition, enrichment of epoxy-eicosatrienoic acid (EET) and dihydroxyeicosatrienoic acid (DHET) synthesis pathways suggests involvement of arachidonic acid-derived lipid mediators that regulate inflammation, vascular tone, and metabolic signaling [18, 19]. These pathways provide a plausible mechanistic bridge between sleep-dependent inflammatory regulation and diabetes risk.

Finally, regional association plots generated by LocusZoom.js [15] demonstrate localized interaction peaks residing within the LD blocks near the *LAMA2* gene on chromosome 6, *NTN1* gene on chromosome 17, and *GRHL1* gene on chromosome 2 (Figure S46 C-E). Together, these functional, pathway-level, and locus-level validations highlight that GETAP identified biologically insightful G↑sleep interaction loci.

### 4.7 Conclusions from real data analyses

Across different phenotype-environment combinations spanning glycemic regulation, pulmonary function, adiposity, and metabolic and inflammatory biomarkers, the proposed GETAP procedure demonstrated robustly powerful performance in the large-scale UK Biobank data analyses. These analyses considered different types of environmental/lifestyle factors. The independent GxE loci that we obtained from these analyses are provided in the supplementary materials.

For phenotypes with strong exposure effects, such as HbA1c level and pulmonary phenotypes in relation to cumulative smoking exposure, GETAP identified the highest or similar number of significant GxE interaction signals compared to the other approaches, including the 2df genotypic test (Table S2). In these settings, interaction effects reflected a mixture of additive, dominant, and recessive architectures, and aggregation across the different genotype encodings by GETAP enabled recovery of GxE loci missed by individual single genetic models. For behavioral and dietary exposures, where we found interaction effects to be weaker, GETAP yielded modest but consistent gains over single genetic models and 2df genotypic tests. This was evident for BMI, CRP, and triglycerides paired with dietary or alcohol-related lifestyle factors. These represent limited and sparse GxE signals (Table S2).

Overall, these results highlight that no single genotype encoding uniformly maximizes power across settings. By aggregating evidence across genetic models, GETAP provides a flexible and powerful approach for large-scale testing of gene-environment interactions.

## 5 Discussion

A major challenge in G×E analysis is accounting for genetic model misspecification, a long-recognized issue in genome-wide studies [4, 20, 21]. In practice, most genome-wide interaction analyses default to additive genotype coding. While additive tests are optimal when the true genetic mode of inheritance is additive, they can suffer substantial power loss when the underlying genetic model follows a dominant or recessive pattern. In this work, we propose and systematically evaluate GETAP, a robust framework for genome-wide gene-environment (G×E) interaction analysis. GETAP combines evidence from additive, dominant, and recessive genotype coding-based GxE tests using the Cauchy combination of p-values, yielding a single omnibus test that is robust to uncertainty in the underlying genetic inheritance model. Through extensive simulation studies and nine large-scale real-data analyses in the UK Biobank, we demonstrate that GETAP provides well-calibrated inference and competitive statistical power across various G↑E settings. GETAP offers greater power than 1df single genetic model-based tests in many settings. The additive genetic model appeared to be the most robust and powerful among the 1df tests. GETAP performs competitively with the additive genetic model in terms of power when the true genetic model is additive or dominant. GETAP outperforms the additive model when the true genetic model is recessive.

In the simulations, we compared GETAP with the additive, dominant, and recessive models individually. Therefore, the comparison corresponds to the scenario in which we choose a single genetic model, such as dominant, to implement genome-wide GxE testing, versus the scenario in which we employ GETAP for whole-genome-wide analysis. The correctly specified single-model test consistently achieved the highest power, as expected. However, GETAP closely tracked the performance of this oracle test while performing substantially better than the misspecified single-model tests. We also notice in simulations that GETAP is particularly advantageous over the additive test when the true genetic mode of inheritance is recessive. We observe the same pattern in the real data applications as well. When the recessive test identifies a substantial number of GxE signals for a phenotype-environment combination, GETAP detects substantially more GxE signals than others, including the recessive model. Across all the phenotype-environment combinations analyzed in the UKB, GETAP performed competitively and identified more GxE loci for most.

Several robust test statistics have been proposed to address model misspecification in genetic association studies [7, 8, 9]. One popular approach is to take the maximum of test statistics across multiple genetic models. For example, the MAX3 or So-Sham test statistic [9] considers the maximum of the three test statistics obtained from the additive, recessive, and dominant model-based tests. The overall p-value is calculated while appropriately adjusting for multiple comparisons. Under the null hypothesis of no interaction, additive, dominant, and recessive G×E test statistics are correlated because they are computed from the same genotype-phenotype-environment data. As a result, max-type approaches, e.g., MAX3, require evaluation of the joint null distribution of correlated statistics, which depends on their covariance structure [22, 8, 9]. Consequently, accurate calibration of MAX-type p-values usually requires multivariate normal tail probability evaluation or carefully designed approximations. Although statistically valid, these procedures introduce additional analytical and computational challenges, particularly in genome-wide scans involving hundreds of thousands of variants. Moreover, in regression-based G×E analyses with covariate adjustment, the covariance structure can vary across variants due to differences in minor allele frequency, genotype distribution, etc.

In contrast, the Cauchy aggregation operates directly on marginal p-values and accounts for arbitrary dependence between the tests. Crucially, it is computationally very fast. This simplicity makes the Cauchy combination of p-values an attractive alternative for a robust GxE testing procedure in large datasets. It can be implemented as a lightweight post-processing step on top of existing regression-based pipelines, such as those implemented in PLINK [23]. Consequently, it scales to a million variants and multiple phenotype-environment combinations naturally. This simplicity is particularly valuable for large-scale G×E studies in modern biobanks, where computational efficiency is a critical consideration.

We also performed a detailed comparison with the genotypic 2df interaction test. GETAP performs better than the 2df test when the actual genetic model is additive or dominant. Otherwise, the power performance is comparable. The 2df test is a genetic-model-free alternative because it does not assume a specific genotype-coding scheme. However, this robustness comes at the cost of an additional degree of freedom, which can reduce power when the true genetic mode of inheritance is simple and particular, e.g., additive or dominant. Previous work on 2df G×E tests has primarily focused on power and sample size calculations rather than genome-wide applications [7]. Our study, to our knowledge, provides one of the first systematic genome-wide comparisons of 1df and 2df interaction tests in large-scale biobank data. In real data analyses, GETAP identified more G↑E signals than the 2df test for most of the phenotype-environment combinations.

The UK Biobank applications highlight the practical utility of GETAP in real-world settings characterized by heterogeneous genetic architectures, diverse phenotype distributions, and various environmental exposures. Across phenotypes related to glycemic regulation, pulmonary function, adiposity, inflammation, metabolism, and cardiometabolic disease, GETAP consistently identified a similar or larger number of GxE loci than those from single-model or 2df tests. For several phenotype-environment pairs, such as HbA1c or pulmonary function in interaction with cumulative smoking exposure, GETAP identified a substantially large number of GxE interaction signals. In the analysis of type 2 diabetes as a binary disease phenotype and sleep duration as an environmental factor, GETAP identified 563 independent GxE loci, which is highly significant compared to the existing literature on GxE studies. For dietary and behavioral lifestyle factors, including alcohol intake frequency and healthy diet score, interaction signals were more diffuse, consistent with prior observations in G↑E studies of complex phenotypes [21]. Even for these phenotype-environment combinations with limited GxE signals, GETAP yielded a modest but consistently higher number of signals than the other approaches.

Throughout this study, we adjusted for multiple testing using the Benjamini-Hochberg (BH) false discovery rate (FDR) controlling procedure [24]. G×E interaction effects are typically minor and sparse. The primary goal of our work is to identify a set of promising GxE loci for downstream investigation rather than to control the stringent family-wise error rate (FWER). In this context, FDR control offers a desirable balance between discovery and error control. FWER control is more relevant for detecting main genetic effects on complex phenotypes, as demonstrated by GWAS. The BH procedure remains valid under the positive dependence structures commonly observed among SNP-level test statistics due to linkage disequilibrium [25]. More conservative FDR controlling procedures would substantially reduce power without much practical benefit. The QQ plots, genomic inflation factor estimation, and sample-size-standardized genomic inflation factors further support the conclusion that the discovered GxE loci are primarily driven by polygenic GxE effects, not by the lack of covariate adjustment or confounding [26].

We conducted functional annotation and pathway analyses for selected phenotype-environment pairs to assess biological relevance of the GxE SNPs using an established post-GWAS tool FUMA [14]. For both HbA1c ↑ smoking and T2D ↑ sleep duration, GxE interaction loci identified by GETAP were enriched in noncoding and regulatory genomic regions. Gene-based and pathway analyses highlight biologically relevant processes related to metabolism, detoxification, inflammation, and lipid signaling. These analyses are exploratory and do not focus on causality, but they reassure that the GxE loci identified by GETAP are biologically meaningful.

We next discuss some limitations of our work. First, GETAP aggregates only three standard genotype encodings. Real genetic architectures may involve additional complex patterns, such as overdominance [20, 21]. GETAP can be extended to aggregate distinct additional models as well. Second, recessive model-based tests can exhibit inflated type I error rates for rare variants and binary phenotypes, and this inflation may propagate to the aggregated p-value. However, large sample sizes of biobank datasets can alleviate this limitation. We need to consider the MAF threshold in the analysis to ensure there is an adequate number of individuals in the rare homozygous genotypic group. While GETAP generally showed improved calibration relative to recessive and 2df tests in our analyses, careful quality control is important. Third, we restricted the analyses in the UK Biobank to individuals of White British ancestry. We plan to extend these analyses to people of diverse ancestries, such as South Asians. Fourth, due to limited computing resources, we were unable to implement the 2df genotypic tests for binary disease phenotypes on the UKB cloud computing platform. Once more resources become available, we will perform these analyses. Finally, as with most G×E studies, measurement errors in environmental exposures may attenuate estimates of interaction effects.

In summary, GETAP provides a simple, scalable, and well-calibrated approach for genome-wide G×E analysis under genetic model uncertainty. By aggregating additive, dominant, and recessive genetic model-based tests of interaction using p-value combination, it offers a robust and powerful approach to G×E testing. As biobank-scale data resources continue to expand, GETAP can be applied to many phenotype-environment combinations and better characterize their gene-environment interplay.

## Data availability

The data used in the study are either simulated or belong to the UK Biobank resource. The real data are available from the UK Biobank. Restrictions apply to the availability of these data, which were used under license for the current study (Application Number 77327) and are not publicly available.

Supplementary materials, analysis scripts, and processed data supporting this study have been deposited in the *GSA Figshare* repository and will be made publicly available upon publication.

## Code availability

Genome-wide gene–environment (G↑E) interaction analyses were conducted using PLINK (v1.9) under additive, dominant, and recessive genotype encodings. The proposed GETAP procedure and all down-stream analyses were implemented in R (Supplementary Data). Variant-level interaction *p*-values were aggregated using ACAT R package.

## Acknowledgement

This research has been conducted using the UK Biobank Resource under Application Number 77327.

## References

[1] R Ottman. Gene-environment interaction: definitions and study design. Preventive medicine, 25(6):764–770, 1996.

[2] Peter Kraft, Yu-Chung Yen, Daniel O Stram, Joyce Morrison, and W James Gauderman. Exploiting gene-environment interaction to detect genetic associations. Human heredity, 63(2):111–119, 2007.

[3] Duncan Thomas. Gene-environment-wide association studies: emerging approaches. Nature reviews genetics, 11(4):259–272, 2010.

[4] Amadou Gaye and Scott K Davis. Genetic model misspecification in genetic association studies. BMC research notes, 10(1):1–9, 2017.

[5] Ali Pazokitoroudi, Alec M Chiu, Kathryn S Burch, Bogdan Pasaniuc, and Sriram Sankararaman. Quantifying the contribution of dominance deviation effects to complex trait variation in biobank-scale data. The American Journal of Human Genetics, 108(5):799–808, 2021.

[6] Matteo Di Scipio, Mohammad Khan, Shihong Mao, Michael Chong, Conor Judge, Nazia Pathan, Nicolas Perrot, Walter Nelson, Ricky Lali, Shuang Di, et al. A versatile, fast and unbiased method for estimation of gene-by-environment interaction effects on biobank-scale datasets. Nature Communications, 14(1):5196, 2023.

[7] Catherine M Moore, Seth A Jacobson, and Tasha E Fingerlin. Power and sample size calculations for genetic association studies in the presence of genetic model misspecification. Human heredity, 84(6):256–271, 2020.

[8] Hon-Cheong So and Pak C Sham. Robust association tests under different genetic models, allowing for binary or quantitative traits and covariates. Behavior genetics, 41(5):768–775, 2011.

[9] Zhongxue Chen and Yong Zang. Cmax3: A robust statistical test for genetic association accounting for covariates. Genes, 12(11):1723, 2021.

[10] Daniel J Wilson. The harmonic mean p-value for combining dependent tests. Proceedings of the National Academy of Sciences, 116(4):1195–1200, 2019.

[11] Yaowu Liu, Shengrui Chen, Zilin Li, Alanna C Morrison, Eric Boerwinkle, and Xiang Lin. Acat: a fast and powerful p-value combination method for rare-variant analysis in sequencing studies. The American Journal of Human Genetics, 104(3):410–421, 2019.

[12] Yaowu Liu and Jun Xie. Cauchy combination test: a powerful test with analytic p-value calculation under arbitrary dependency structures. Journal of the American Statistical Association, 115(529):393– 402, 2020.

[13] Paul IW De Bakker, Manuel AR Ferreira, Xiaoming Jia, Benjamin M Neale, Soumya Raychaudhuri, and Benjamin F Voight. Practical aspects of imputation-driven meta-analysis of genome-wide association studies. Human molecular genetics, 17(R2):R122–R128, 2008.

[14] Kyoko Watanabe, Erdogan Taskesen, Arjen Van Bochoven, and Danielle Posthuma. Functional mapping and annotation of genetic associations with fuma. Nature communications, 8(1):1826, 2017.

[15] Andrew P Boughton, Ryan P Welch, Matthew Flickinger, Peter VandeHaar, Daniel Taliun, Gonçalo R Abecasis, and Michael Boehnke. Locuszoom. js: interactive and embeddable visualization of genetic association study results. Bioinformatics, 37(18):3017–3018, 2021.

[16] Daniel W Nebert and Timothy P Dalton. The role of cytochrome p450 enzymes in endogenous signalling pathways and environmental carcinogenesis. Nature reviews cancer, 6(12):947–960, 2006.

[17] Laila AlDabal and Ahmed S BaHammam. Metabolic, endocrine, and immune consequences of sleep deprivation. The open respiratory medicine journal, 5:31, 2011.

[18] Koichi Node, Yuqing Huo, Xiulu Ruan, Baichun Yang, Martin Spiecker, Klaus Ley, Darryl C Zeldin, and James K Liao. Anti-inflammatory properties of cytochrome p450 epoxygenase-derived eicosanoids. Science, 285(5431):1276–1279, 1999.

[19] Zeqi Shi, Zuowen He, and Dao Wen Wang. Cyp450 epoxygenase metabolites, epoxyeicosatrienoic acids, as novel anti-inflammatory mediators. Molecules, 27(12):3873, 2022.

[20] David J Hunter. Gene-environment interactions in human diseases. Nature reviews genetics, 6(4):287– 298, 2005.

[21] Duncan Thomas. Methods for investigating gene-environment interactions in candidate pathway and genome-wide association studies. Annual review of public health, 31(1):21–36, 2010.

[22] B Freidlin, G Zheng, Z Li, and J L Gastwirth. Trend tests for case-control studies of genetic markers: power, sample size and robustness. Human heredity, 53(3):146–152, 2002.

[23] Shaun Purcell, Benjamin Neale, Kathe Todd-Brown, Lori Thomas, Manuel AR Ferreira, David Bender, Julian Maller, Pamela Sklar, Paul IW De Bakker, Mark J Daly, et al. Plink: a tool set for whole-genome association and population-based linkage analyses. The American journal of human genetics, 81(3):559–575, 2007.

[24] Yoav Benjamini and Yosef Hochberg. Controlling the false discovery rate: a practical and powerful approach to multiple testing. Journal of the Royal Statistical Society: Series B (Methodological), 57(1):289–300, 1995.

[25] Yoav Benjamini and Daniel Yekutieli. The control of the false discovery rate in multiple testing under dependency. Annals of statistics, pages 1165–1188, 2001.

[26] Jian Yang, Michael N Weedon, Shaun Purcell, Guillaume Lettre, Karol Estrada, Cristen J Willer, Albert V Smith, Erik Ingelsson, Jeffrey R O’connell, Massimo Mangino, et al. Genomic inflation factors under polygenic inheritance. European journal of human genetics, 19(7):807–812, 2011.

[27] Emelia J Benjamin, Michael J Blaha, Stephanie E Chiuve, Mary Cushman, Sandeep R Das, Rajat Deo, Sarah D De Ferranti, James Floyd, Myriam Fornage, Cathleen Gillespie, et al. Heart disease and stroke statistics—2017 update: a report from the american heart association. circulation, 135(10):e146–e603, 2017.

